# Bursty Gene Expression in Single Cells and Expanding Populations: A Discrete Approach

**DOI:** 10.1101/2025.05.20.655093

**Authors:** Jakub Poljovka, Iryna Zabaikina, Pavol Bokes, Abhyudai Singh

**Author notes:** {, }.

## Abstract

Bursty protein production is a key source of gene expression noise. In this study, we analyze a Markovian model of cellular protein dynamics, where proteins are produced in geometrically distributed bursts. Extending this model, we incorporate a feedback mechanism by assuming that higher protein levels reduce both cell growth and protein decay rates. We study both single-cell dynamics and an expanding cell population. Without feedback, the protein level follows a negative binomial distribution with the same parameters in both cases. With feedback, however, single-cell and population-level distributions differ, each expressible as a mixture of two negative binomial distributions with framework-dependent parameters. Using numerical integration of the master and population balance equations, we calculate the time-dependent distributions in both settings. This work extends previous continuous models and provides new insights into how population expansion influences intrinsic cellular heterogeneity.

## I INTRODUCTION

Because biochemical reactions are random in nature, gene expression is inherently stochastic [1]–[4]. A significant portion of this variability arises from the synthesis of gene products in bursts, where multiple molecular copies are produced in rapid and brief transcription/translation events, followed by longer periods of quiescence [5]–[13]. Gene expression variability (noise) can be modeled using stochastic processes, particularly Markov models [14], [15]. We focus on a class of models in which production bursts of a given protein are treated as instantaneous events that occur randomly in time, increasing the protein level by a value known as the burst size [16]–[18].

The protein level can be modeled as a discrete or continuous variable. In continuous models, the burst size is typically assumed to follow an exponentially distributed random variable [19], whereas in discrete models, it is often modeled by a geometric distribution [20], [21]. Protein production is balanced by decay, which may result from active degradation or dilution due to cell growth. In continuous models, decay is typically described using deterministic ordinary differential equations [22]. In discrete models, decay is represented by transitions that decrease the protein level by one unit [23].

An important feature of gene expression is feedback regulation, where the protein itself modulates its own abundance [24]–[26]. We focus here on positive feedback on protein decay, in which increased protein levels reduce the decay rate [27], [28]. We specifically examine the case where the decay rate is tied to the cell growth rate, a scenario that occurs when decay is driven by the dilution effect of cell volume expansion [29], [30]. Positive feedback resulting from such expression-growth coupling have been reported across cell types, where stochastic expression of certain gene products reduces cellular growth but allows bet hedging against uncertain changes in the extracellular environment [31]–[34].

Previous studies have explored this feedback within continuous protein models, reporting that the probability distribution of protein levels in a single cell differs from the distribution observed in a population of growing cells [35]–[37]. In particular, the probability of residing in a saturated protein state is overestimated in the single-cell view compared with the population-wide view. In this study, we investigate gene expression using a discrete protein model. Our approach not only reinforces previous findings but also extends the methodology to discrete systems, providing new explicit stationary distributions. Moreover, we develop a framework for numerical integration in the discrete model, allowing us to compute time-dependent distributions for both individual cells and growing cell populations.

## II. Modeling stochastic gene expression

We model the dynamics of the protein level in a single cell using a continuous-time Markov process *n*(*t*), where the state space consists of nonnegative integers. The variable *n*(*t*) represents a quantized protein concentration — specifically, the integer part of the protein concentration per unit volume at time *t*. Consequently, an increase in cell volume leads to a reduction in *n*(*t*), a phenomenon we refer to as *dilution* (Fig. 1).

**Fig. 1.**
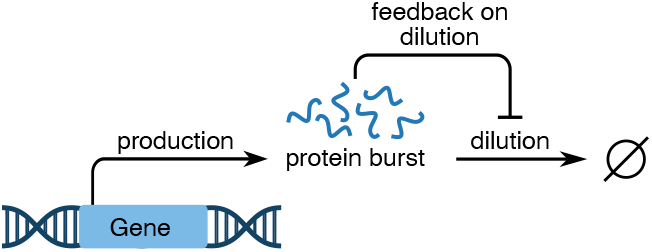
Schematic of the gene expression model. Protein is produced in random bursts, and between bursts, its concentration decreases due to dilution. The dilution rate depends on the protein level, introducing a feedback mechanism.

The Markov process is specified by its transition intensities. In this model, we assume that within an infinitesimal interval *dt*, a protein burst occurs with probability *λdt*, where *λ* is the burst frequency. Each burst event increases the protein level by *b* units; the burst size *b* follows a geometric distribution, *b* ∼ Geom(*β*), with the mean burst size *β*. Simultaneously, protein is lost due to dilution with a rate *f* (*n*), so the probability that the protein level decreases by one within *dt* is *f* (*n*)*dt*. The dilution rate *f* (*n*) is obtained by multiplying the cell growth rate *r*(*n*) by the protein level *n*:

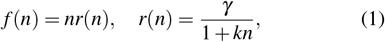

where the parameter *k* quantifies the strength of the feedback; specifically, when *k* = 0, the growth rate is constant and the dilution rate is proportional to the protein level: there is no feedback effect. On the other hand, positive values of *k* introduce a positive feedback mechanism: as the protein level increases, the cell growth slows down, and so does dilution, thus preserving the protein level. These two reactions – bursty production and dilution – constitute possible transitions in the single-cell Markov model.

To account for a growing cell population, we incorporate an additional process: cell proliferation. We assume that each cell divides at a protein-dependent rate denoted by *r*(*n*), so that the probability of division within an infinitesimal time interval *dt* is *r*(*n*)*dt*. Upon division, the cell volume is halved and the protein molecules are assumed to be equally partitioned between the two daughter cells. As a result, the protein level *n* in each daughter cell remains unchanged. A summary of both the single-cell dynamics and population-level processes is provided in Figs. 2 and 3.

**Fig. 2.**
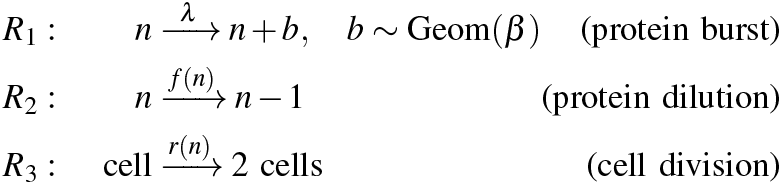
Scheme of the stochastic process. Reactions *R*_1_ and *R*_2_ specify the Markovian transitions between protein levels within one cell. Reaction intensities are detailed on top of the reaction arrows. Reaction *R*_3_ creates a replica of the process — a daughter cell. The new cell behaves independently of its mother. The single-cell framework omits reaction *R*_3_.

**Fig. 3.**
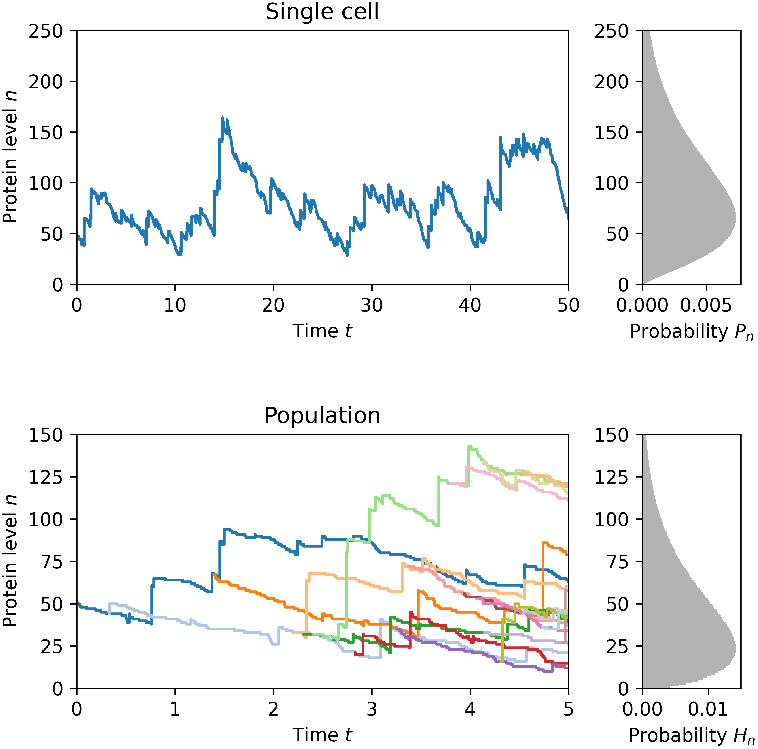
Single-cell and population processes. The simulation is initiated at *t* = 0 with a single cell with protein level *n* = 50 and follows the reaction scheme in Fig 2. The right-hand panels show large-time protein-level distributions *P*_*n*_ and *H*_*n*_ for the two formulations, which are generally different, and for which we derive explicit formulae in this work. As for parameter values, we use burst frequency *λ* = 2.33, mean burst size *β* = 10 and dilution/cell-growth rate (1) with *γ* = 1 and *k* = 0.03.

## III. Expression heterogeneity in single-cell perspective

In this section, we formulate the master equation and the associated steady-state equation for a single-cell model. We show that the generating function of the steady-state probability distribution satisfies a linear first-order ordinary differential equation. By solving this equation and reverting the generation function back to the probability function space, we obtain an explicit steady-state distribution. The variables and parameters introduced in this section, as well as throughout the paper, are summarized in Table I.

**TABLE I.**
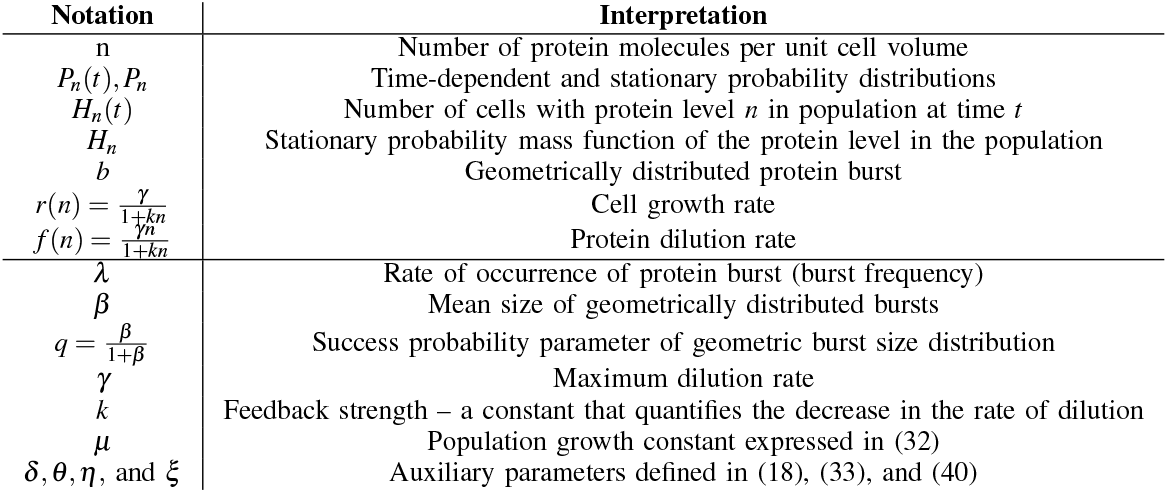
Model parameters and variables studied in the text.

Let *P*_*n*_(*t*) denote the probability that the protein level is *n* at time *t*. Under the assumptions stated in Section II, it satisfies the following master equation:

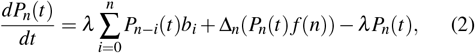

where Δ_*n*_(·) is the forward difference operator. On the right-hand side of (2), the first term is the influx of probability into the state *n* due to burst production (including absence of the burst as *i* = 0), the second term is the net flux of probability into state *n* due to dilution, and the third term is the outflux of probability from *n* caused by bursts. The probability mass function of the burst size *b*_*i*_ represents the probability of producing *i* proteins, which is given by:

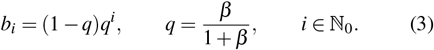

Let *P*_*n*_ denote the probability that the protein level is *n* in the stationary state. In steady state, the probability distribution *P*_*n*_ is independent of time, which also implies that *dP*_*n*_(*t*)*/dt* = 0. Then, the master equation (2) simplifies to the following equation for the stationary probability distribution:

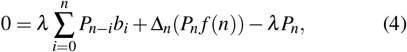

which can be stated in the equivalent form:

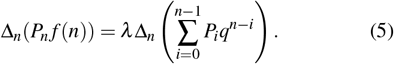

By applying antidifferencing to equation (5), we obtain:

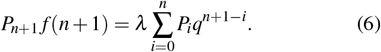

Next, we substitute *f* (*n*) defined in (1) into (6) yields:

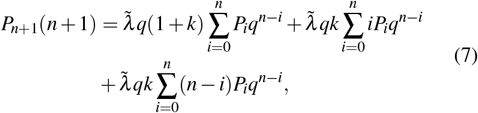

where the parameter 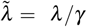 is the burst frequency normalized by the maximum dilution rate.

For further analysis, we introduce the (probability) generating function *G*_*Y*_ (*x*) of the discrete random non-negative variable *Y* :

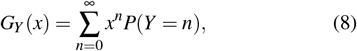

which is analytic, converges absolutely for all *x* ∈ [0, 1], and satisfies *G*_*Y*_ (1) = 1. We additionally require it to be positive; it ensures that the corresponding probability distribution is well-defined.

The generating functions allows us to rewrite the convolution in summations as the product of the generating functions:

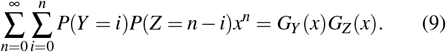

Let us define the generating function *G*_*P*_(*x*) of the distribution *P*_*n*_:

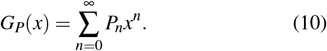

Since the right-hand side of equation (7) consists of three convolutions, we employ the convolution property (9) and obtain the following differential equation for *G*_*P*_(*x*):

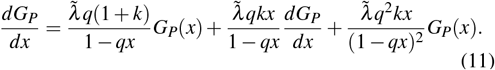

We obtain a solution of (11) in the form:

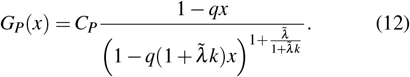

However, the solution (12) must satisfy the conditions of a valid generating function as given in (8). In particular, its denominator must remain positive for all *x* ∈ [0, 1]; otherwise *G*_*P*_(*x*) becomes singular. Since the denominator has minimum at *x* = 1, the following condition on parameters must hold:

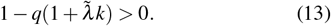

Substituting, *x* = 0 into (12), we find that *C*_*P*_ = *P*_0_. Next, substituting *x* = 1 into (12), we obtain:

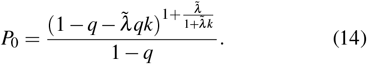

Expanding (12) into a power series in *x* yields:

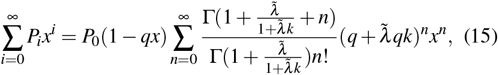

where Γ(·) denotes the gamma function.

Comparing terms with the same powers of *x* in (15) yields the stationary probability mass function of the protein level in the single cell:

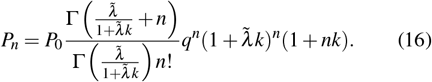

Note that is also possible to express *P*_*n*_ as a weighted average of two negative binomial mass functions:

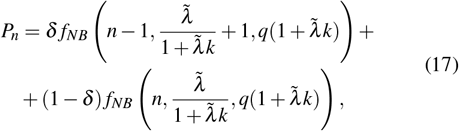

where

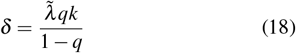

and *f*_*NB*_(*n, r, p*) represents the mass function of the negative binomial distribution, given by:

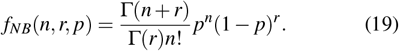

Note that in absence of feedback special (*k* = 0), (17) takes the form of a negative binomial distribution, this is a well-known result [20].

Finally, by utilizing the fact that 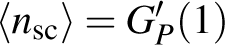, we obtain the explicit expression of the mean protein level in the single cell (sc):

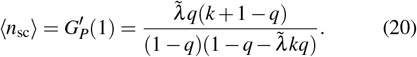

For *k* = 0, the mean protein level ⟨*n*_sc_⟩ matches that in negative binomial distribution; for *k >* 0, ⟨*n*_sc_⟩ deviates from the negative binomial prediction, reflecting the impact of growth-dependent dilution. The effect of increasing *k* on the protein distribution is visualised in Fig. 4.

**Fig. 4.**
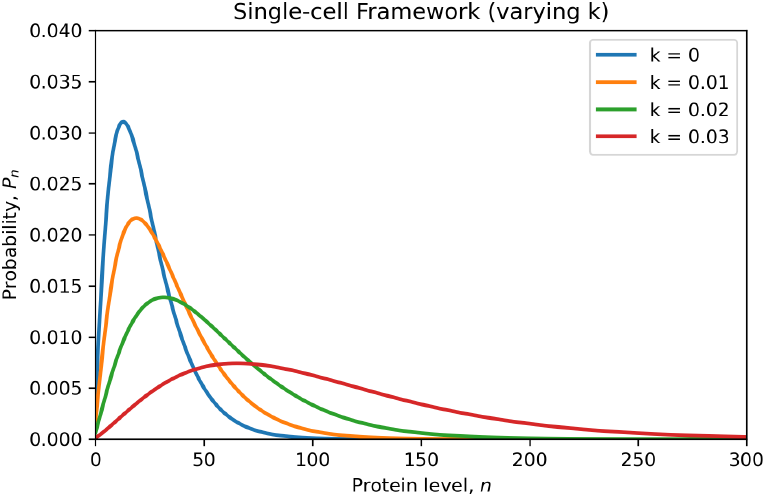
The effect of increasing the positive feedback strength on the protein distribution in single cells.

## IV. Expression heterogeneity in population-level perspective

In this section, we formulate a population balance equation, which describes the temporal evolution of the population-wide protein distributions. Like in the single-cell case, we transform the equation into the generating function space. An additional subtlety is here posed by the unknown principal eigenvalue (the Malthusian parameter) for the asymptotic population growth. We isolate the eigenvalue, and its corresponding eigenvector, by considering regularity conditions for the generating function.

Let *H*_*n*_(*t*) denote the number of cells in population with the protein level *n* at time *t*. Then the time evolution of *H*_*n*_(*t*) is governed by the following population balance equation:

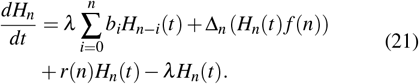

Apart from the change in the symbol for the dependent variable, the population balance equation (21) differs from the master equation (2) only by the inclusion of the growth term *r*(*n*)*H*_*n*_(*t*). We apply the separation of variables and look for the solution to (21) in the form

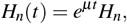

where *µ* is the principal eigenvalue (the one with the largest real part) and *H*_*n*_ is the principal eigenvector of (21). The principal exponential mode is asymptotically dominant as *t*→ ∞ for solutions of (21), thus providing their large time behavior. Then *µ* represents the growth rate of the population (the Malthusian parameter) and *H*_*n*_ is the stationary protein distribution in the cell population. Substituting *H*_*n*_(*t*) = *e*^*µt*^*H*_*n*_ into (21), we obtain:

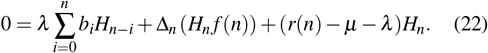

Let us define an auxiliary sequence:

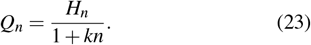

Let us denote *G*_*H*_(*x*) and *G*_*Q*_(*x*) as the generating functions of *H*_*n*_ and *Q*_*n*_, respectively:

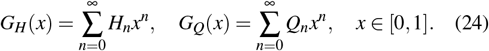

By using the convolution property (9), we obtain the following equation:

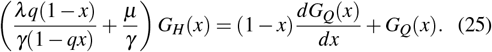

Through direct substitution of (23) into (24), we obtain that *G*_*H*_(*x*) can be expressed in terms of *G*_*Q*_(*x*) as follows:

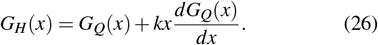

Next, we substitute (26) into (25) and obtain:

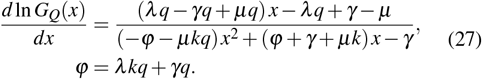

Next, we perform a partial fractions decomposition of the right-hand side using. We define the discriminant of the quadratic in the denominator as follows:

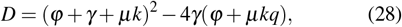

which is positive for all permissible parameter values. The roots of the denominator in (27) are given by:

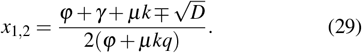

Rewriting equation (27) in terms of partial fractions yields:

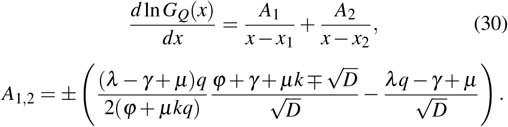

Integrating and exponentiating both sides of equation (30), we obtain the solution:

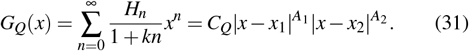

It can be shown that *x*_1_ *<* 1 and *x*_2_ *>* 1. Then, assuming *A*_1_ ≠ 0, we find that *G*_*Q*_(*x*_1_) is either zero or singular at *x* = *x*_1_, contradicting the definition of the generating function (8). Therefore, we set *A*_1_ = 0 and obtain the value of the Malthusian parameter *µ*:

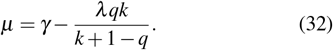

Since *µ* is also the population growth rate, we require that *µ* is positive. Otherwise, the population would extinct.

The term *A*_2_ from (30) simplifies using the fact that *A*_1_ = 0; it can also be shown that *A*_2_ *<* 0. Thus, we introduce an additional parameter *η*, defined as its opposite value, and define a new parameter *θ* as the reciprocal of *x*_2_:

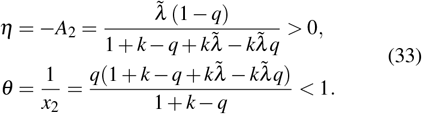

Next, we remove the absolute value in (31) by leveraging the condition *x*_2_ *>* 1. As the result, the generating function *G*_*Q*_(*x*) in (31) simplifies as follows:

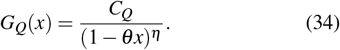

The normalization condition for the probability mass function *H*_*n*_ can be expressed in terms of the generating function as 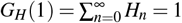. Furthermore, by substituting *x* = 1 into (25), we obtain *G*_*Q*_(1) = *µ/γ*; therefore we can express the constant as follows:

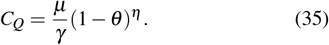

Expanding the right-hand side of (34) into a power series in *x* yields:

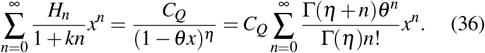

By comparing the terms with the same powers of *x*, we obtain the stationary probability mass function of the protein level in the cell population:

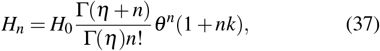

where:

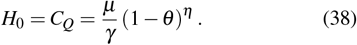

Note that similar to the single-cell model, *H*_*n*_ in (37) is also a weighted average of two negative binomial mass functions:

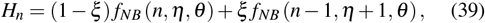

where

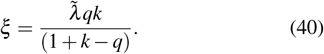

The population mean of protein level ⟨*n*_pop_⟩ is calculated as 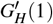 and given by:

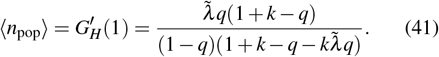

Note that for *k* = 0, both the single-cell and population-level models share the same mean and distribution, which is consistent with previous studies [29], [35]. The effect of increasing the feedback strength *k* is visualised in Fig. 5

**Fig. 5.**
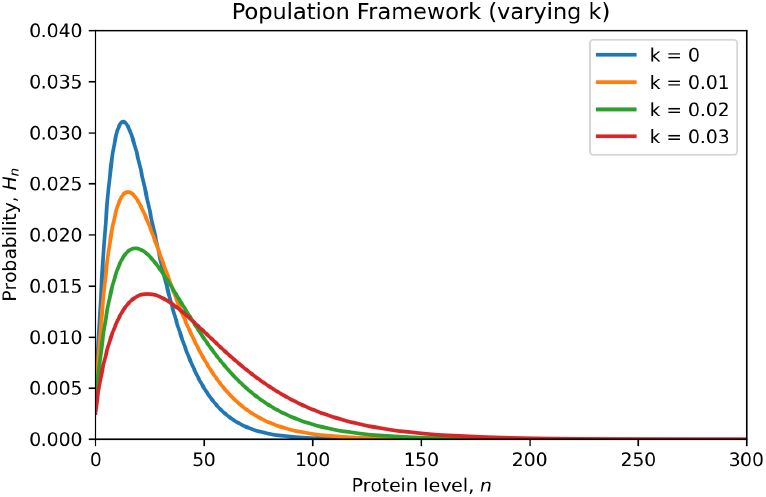
The effect of increasing the positive feedback strength on the protein distribution in growing populations.

## V. Numerical approach to solving master equations

In general, the master equations (2) and (21) can also be represented in matrix form as:

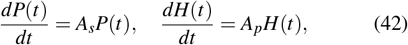

where *P*(*t*) is probability vector describing the protein level in the single-cell model and *H*(*t*) is vector describing the number of cells with given protein level in the population model. The matrices *A*_*s*_ and *A*_*p*_ correspond to the single-cell and population models, respectively, and have the following general form:

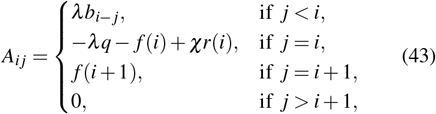

where *χ* is the indicator parameter such that *χ* = 0 in *A*_*s*_ and *χ* = 1 in *A*_*p*_. For the equations (42), the solutions can be expressed in the following form:

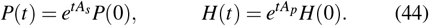

The main challenge in the numerical solution is that the vectors *P*(*t*) and *H*(*t*), as well as the matrices *A*_*s*_ and *A*_*p*_, are infinite-dimensional. Thus, for numerical computations, we must truncate the dimension at a finite level *N*, which represents the maximum protein concentration [38], [39]. This numerical approximation allows us to solve the master equations (2) and (21) for any arbitrary initial distributions. The results are shown in Fig. 6, which illustrates the dynamics of the protein distribution in the single-cell (Fig. 6A) and population (Fig. 6B) frameworks, respectively. In both cases, we observe that the process converges from its initial state *t* = 0 to its steady state as *t* → ∞. The convergence is more rapid in the population framework. The population distribution is shifted to the left compared to the single cell distribution, which is a consequence of the dilution-coupled feedback, as cells with higher protein level divide less frequently.

**Fig. 6.**
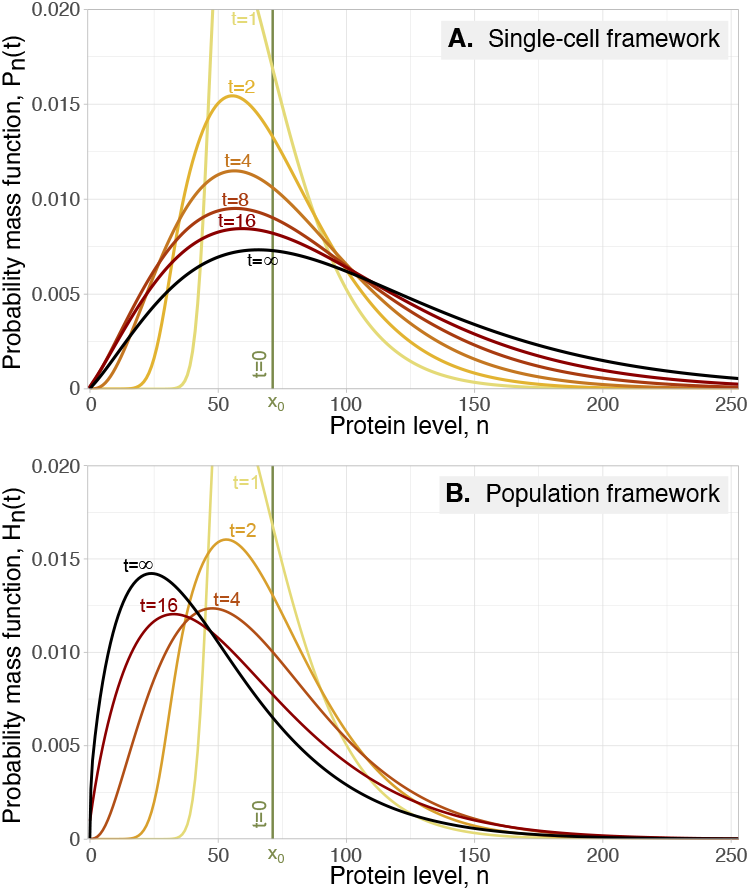
Time evolution of protein distributions in single-cell and population-level frameworks. **A:** The time-dependent single-cell distributions *P*_*n*_(*t*) (colored lines) evolve from the initial condition *P*_*n*_(0) and converge as *t* → ∞ to the stationary distribution *P*_*n*_ (black). **B:** Similarly, the population-level distributions *H*_*n*_(*t*) (colored lines) evolve from the initial condition *H*_*n*_(0) and approach as *t* → ∞ the corresponding stationary distribution *H*_*n*_ (black). In both cases, the initial condition is set to a delta distribution *P*_*n*_(0) = *H*_*n*_(0) = *δ*_*n*,70_. A numerical cut-off *N* = 600 for the protein level is used. Model parameters are: *λ* = 2.33, *β* = 10, *k* = 0.03, and *γ* = 1.

## VI. CONCLUSIONS

### A. Results

In this study, we investigate a discrete bursty protein model that incorporates the regulation of protein decay and cell growth. The discrete protein level variable is interpreted as a quantized protein concentration, defined as the number of protein molecules per unit cell volume. A key feature of our model is the quantitative coupling between the protein decay rate and the cell growth rate. This relationship arises because an increase in cell growth leads to an expansion in cellular volume, which in turn results in a greater reduction in protein concentration.

To simplify the analysis, we neglect the effects of partitioning errors at cell division. This approximation is justified by the fact that, while the absolute number of protein molecules is divided between daughter cells, the cell volume is also partitioned. Consequently, the fluctuations introduced by partitioning errors remain relatively small and do not significantly impact the overall protein concentration dynamics.

A key advantage of our simplified model is its analytical tractability at steady state, allowing us to gain valuable insights into the differences between single-cell and population-level frameworks. The single-cell framework describes the probability distribution of protein levels within an individual cell, while the population framework predicts the fraction of cells in an exponentially growing population that express a given protein level. In the absence of protein-mediated regulation of cell growth, both frameworks yield the same classical negative binomial distribution for protein levels. However, we demonstrate that when protein regulation of cell growth is introduced, the single-cell and population-level distributions diverge. In both cases, we show that the resulting distributions can be expressed as probabilistic mixtures of two distinct negative binomial probability mass functions (17), (39).

The time-dependent probability and population distributions can be obtained by solving a system of linear ordinary differential equations. These solutions can be computed using implicit or explicit time discretization methods, as well as matrix exponentiation techniques. As time progresses, the time-dependent distributions asymptotically approach the explicit stationary distributions, confirming the steady-state behavior of the system (Fig. 6).

### B. Future Work

A promising direction for future research is to explore the relationship between discrete and continuous modeling frameworks. In the continuous case, the probability and population balance equations take the form of integro-partial-differential equations [40]. Solving these equations requires discretization of the protein level variable, for instance, using the finite volume method [41]. Within this context, the discrete model presented in this study can be interpreted as a first-order discretization of the continuous model, exhibiting favorable numerical stability properties.

Further extensions of this work include incorporating a broader range of regulatory feedback mechanisms, encompassing both positive and negative feedback loops. Additionally, investigating the combined effects of protein dilution and degradation, as well as addressing optimization problems related to cellular control and resource allocation [30], would provide valuable insights. The methodological framework developed in this study lays a strong foundation for pursuing these research directions.

